# The application of *Nicotiana benthamiana* as a Transient Expression Host to Clone the Coding Sequences of Plant Genes

**DOI:** 10.1101/2022.12.09.519829

**Authors:** Jianzhong Huang, Peng Jia, Xiaoju Zhong, Xiuying Guan, Hongbin Zhang, Honglei Ruan

## Abstract

Coding sequences (CDS) are commonly used for transient gene expression, in yeast two-hybrid screening, to verify protein interactions and in prokaryotic gene expression studies. CDS are most commonly obtained using complementary DNA (cDNA) derived from messenger RNA (mRNA) extracted from plant tissues and generated by reverse transcription. However, some CDS are difficult to acquire through this process as they are expressed at extremely low levels or have specific spatial and/or temporal expression patterns *in vivo*. These challenges require the development of alternative CDS cloning technologies.

In this study, we found that the genomic intron-containing gene coding sequences (gDNA) from *Arabidopsis thaliana, Oryza sativa, Brassica napus*, and *Glycine max* can be correctly transcribed and spliced into mRNA in *Nicotiana benthamiana*. In contrast, gDNAs from *Triticum aestivum* and *Sorghum bicolor* did not function correctly. In transient expression experiments, the target DNA sequence is driven by a constitutive promoter. Theoretically, a sufficient amount of mRNA can be extracted from the *N. benthamiana* leaves, making it conducive to the cloning of CDS target genes. Our data demonstrate that *N. benthamiana* can be used as an effective host for the cloning CDS of plant genes.

## 1. Introduction

The transient expression system of *Nicotiana benthamiana* provides a rapid, high-yield, and cost-effective method for the synthesis of proteins and biopharmaceuticals [1][2][3]. Agrobacterium can be used to transport protein expression constructs into plant cells. Only a few DNA fragments can be integrated into the chromosomes of plant cells, and most foreign DNA molecules remain transcriptionally active for several days [4]. Transient gene expression studies have shown that maximum expression occurs within 18 hours (h) to 48 h after inoculation, and can last for 10 days [5]. Transient expression systems are often used for the analysis of proteins including protein subcellular localization studies, co-immunoprecipitation (Co-IP), and bimolecular fluorescence complementation (BiFC) [6][7].

Generally, transient expression constructs consist of a constitutive promoter, the CDS of the target gene, and tags for protein detection. mRNA derived from plant tissues can be synthesized into cDNA by reverse transcription to acquire the CDS of target genes. However, some CDS are difficult to amplify from cDNA for cloning resulting in extremely low expression levels of the target gene or specific spatial and/or temporal expression patterns in plants. Alternative methods to synthesize CDS, such as chemically synthesizing genes or the assembly of exons into CDS can be used [8][9]. Yet the chemical synthesis of genes is relatively expensive, particularly for long CDS. Also, it is difficult to assemble the CDS for genes that contain many exons, and so there is a need for the development of alternative CDS cloning measures.

As gDNA can be easily amplified from plant genomic DNA, and *N. benthamiana* can function to splice RNA into mature mRNA, we aimed to determine if plant gDNAs can be used for transient expression. We demonstrated that gDNAs from *Arabidopsis thaliana, Oryza sativa, Brassica napus*, and *Glycine max* can be correctly transcribed and spliced into mRNAs in *N. benthamiana*, whilst gDNAs from *Triticum aestivum* and *Sorghum bicolor* are not effective. In conclusion, our results suggested that the transient expression system of *N. benthamiana* is a useful tool for the cloning of CDSs of plant genes.

## 2. Materials and methods

### 2.1 Plant materials and growth conditions

*Arabidopsis* and *N. benthamiana* plants were grown in pots with autoclaved vermiculite and watered with Hoagland solution. The plants were grown at 24°C under a 16 h light/8 h dark cycle. The leaves of *Brassica napus, Glycine max, Triticum aestivum* and *Sorghum bicolor* were obtained as gifts from Dr. Xin Yuan (Wuhan University, China).

### 2.2 Genomic DNA extraction and construction of vectors

The CTAB DNA extraction protocol was used to extract genomic DNA. The pUC19 plasmid was modified into a gateway compatible entry vector pUC19. In short, the gDNA sequence was amplified from plant genomic DNA and transferred into the entry vector pUC19 using one-step cloning technology. Then, the entry vector and the expression vector pEarleygate 101 were recombined by the LR enzyme to produce *35S::gDNA-YFP-HA* protein expression structure for subsequent transient expression experiments. The HA tag was used to detect the transiently expressed proteins.

### 2.3 Transient expression system of N. benthamiana

The transient expression of genes in the leaves of *N benthamiana* leaves was carried out as previously reported [10]. The bacteria were transfected into *N. benthamiana* leaves using a 1 ml needle-less syringe. The *N. benthamiana* plants were maintained in the dark for 24 h after infiltration.

### 2.4 mRNA extraction and cDNA synthesis by reverse transcription

Total RNA was extracted from the plant leaves (0.2 g) using Trizol reagent and trichloromethane. The RNA was precipitated with isopropanol and dissolved in RNase-free water. DNA contaminants were removed by treating the RNA solution with RNase-free DNase at 37°C for 30 minutes (min). First-strand cDNA was synthesized from the total RNA using a Revert Aid First Strand cDNA Synthesis Kit (Thermo Scientific) at 42°C for 60 min. The target CDS were amplified from cDNAs for cloning and transient expression. The HA sequence was used as a reverse primer to specifically amplify the CDS sequence of the transient expression gene.

### 2.5 Quantitative real-time PCR

The housekeeping genes *At_Actin2* (*AT3G18780*) in *Arabidopsis, Nb_Actin3* (*Niben101Scf03493g00020*) in *N. benthamiana*, and *Os_Actin* (*AB047313*) in *Oryza sativa* were used as internal controls, respectively. The relative expression levels were calculated using the 2^−ΔΔCT^ method as described previously [11]. All qRT-PCR experiments were carried out in triplicate together with respective controls. A special forward primer was used to distinguish PLDγ3.1 and 131 PLDγ3.2. The primer was located in an extra 90 bp intron (548-637) that was spliced out in PLDγ3.2 compared to the sequence of PLDγ3.1. In qRT-PCR, we first calculated the total relative expression of PLDγ3. The relative expression of PLDγ3.1 was calculated with a special forward primer. The relative expression of PLDγ3.2 was calculated from the difference between the total relative expression of PLDγ3 and the relative expression of PLDγ3.1.

### 2.6 Protein extraction and detection

Protein extraction and western blotting were performed as previously reported [10]. Proteins were detected with anti-HA antibody (Roche, Cat#11867423001).

### 2.7 Confocal laser scanning microscopy

A confocal laser scanning microscopy (Leica SP8) was used to visualize YFP protein expression at 40 h after transfection at a wavelength of 514 nm.

## 3. Results

### 3.1 gDNAs from Arabidopsis thaliana can be correctly transcribed and spliced into mRNA in N. benthamiana

The gDNAs of seven members of the phospholipase D (PLD) family in *Arabidopsis* (PLDα2, PLDβ1, PLDβ2, PLDδ, PLDε, PLDγ1, and PLDγ3) were cloned from genomic DNA extracted from the leaves of *Arabidopsis*. Each gDNA contained 2-9 introns (Table S1). The gDNAs were cloned into the pEarleyGate 101 expression vector, which was controlled by the constitutive expression of the CaMV 35S promoter. The YFP-HA coding sequence was fused to the 3’ end of the gDNA, and so only properly spliced mRNA could produce proteins with YFP-HA.

The expression constructs were transiently expressed in the leaves of *N. benthamiana*. Our results showed that PLDα2, PLDβ1, PLDβ2, PLDγ3, and PLDδ were detected with an anti-HA antibody (Fig. 1A), indicating that the five gDNAs from *Arabidopsis* were precisely spliced into mRNA in *N. benthamiana*. Coincidentally, PLDα2-YFP-HA and PLDδ-YFP-HA were observed as plasma membrane proteins (Figs. 2A, B), which was consistent with the findings from previous reports [12][13].

**Fig. 1.**
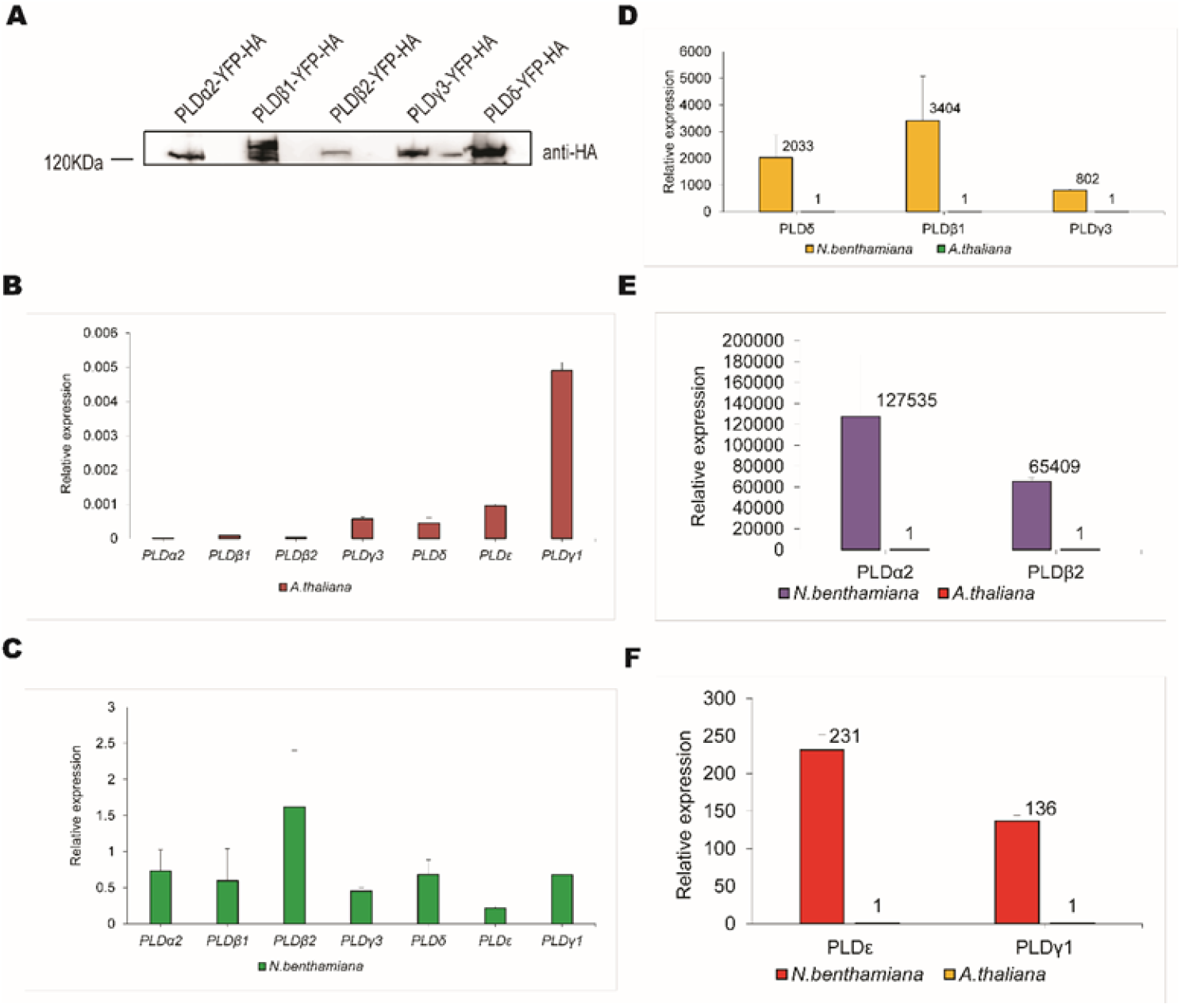
The transient expression of the gDNAs of *AtPLD* genes in the leaves of *N. benthamiana*. (A) The protein expression of YFP-HA tagged PLDα2, PLDβ1, PLDβ2, PLDγ3, and PLDδ. Agrobacteria containing the expression constructs of *35S::gene-YFP-HA* were transfected into the leaves of *N. benthamiana* at OD_600_ of 0.5. The total proteins and mRNAs were extracted and analyzed at 48-hour post infiltration (hpi), and the YFP-HA tagged proteins were detected using an anti-HA antibody. (B-C) qRT-PCR was used to analyze the relative expression of PLD genes in *Arabidopsis* and *N. benthamiana*. The levels of *At_Actin2* (AT3G18780) and *Nb_Actin3* (Niben101Scf03493g00020) were used as internal controls in *Arabidopsis* and *N. benthamiana*, respectively. The values represent the average of three replicate experiments ± SD. (D-F) Comparison of the relative expressions of PLD genes in *N. benthamiana* and *Arabidopsis*. The relative expression level of each gene in *Arabidopsis* was set as 1. The values represented the average of three replicate experiments ± SD.

**Fig. 2.**
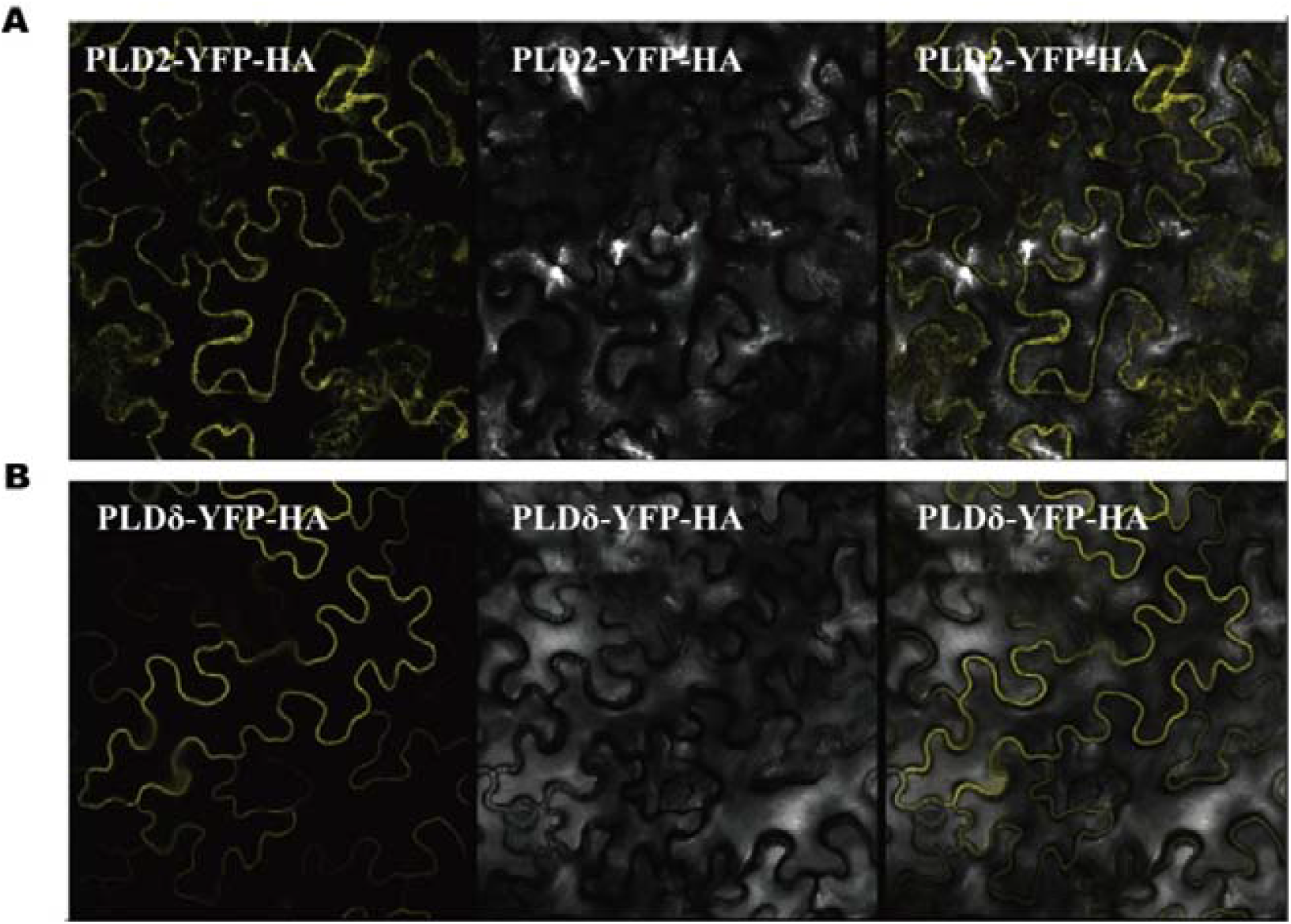
PLDα2 and PLDδ are localized on the plasma membrane. PLDα2 (A) and PLDδ (B) were transiently expressed in the leaves of *N. benthamiana*. Images were pictured at 40 hpi.

The seven PLD family genes had similar high expression levels in *N. benthamiana*, but the protein levels were different (Fig. 1A). PLDγ1 and PLDε was not detected in *N. benthamiana*. The protein level was determined by the rates of synthesis and degradation. Although the mRNA level is a positive indicator for protein synthesis, protein folding, subcellular location, and protein-targeted degradation can affect the stability of different proteins. The different levels of PLD protein members reflects the differential stability of these proteins.

Each cDNA fragment of the transiently expressed PLD gene was amplified and sequenced to determine the splicing of the transient expression gene. Forward primers use the starting part of the PLD gene sequence, while employing the HA sequence inherent in the expression vector as reverse primers to specifically obtain products including the CDS sequence of the PLD gene in *N. benthamiana*. The sequencing results confirmed that all seven PLD gDNAs could form the same mRNAs as in *Arabidopsis*. The protein expression and the mRNA splicing results indicated that the gDNAs from *Arabidopsis* were correctly transcribed and spliced into the mRNA in *N. benthamiana*.

We analyzed the expression of these seven genes in *Arabidopsis* and *N. benthamiana* by qRT-PCR. The seven native PLD genes were differentially expressed in the leaves of *Arabidopsis* (Fig. 1B). There are various factors that affect their expression, including protein generation and degradation, such as the strength of the promoter, the number and position of introns, the growth period of plant, the stimulation of pathogens, and the protein-targeted degradation and so on [5].The expression level of PLDα2 was folds lower than PLDγ1. Although PLDγ1 was expressed at the highest level of the seven genes, the relative expression level of PLDγ1 compared to the control gene *At_Actin2* was 0.0049. Moreover, the transient expression levels of the seven genes in *N. benthamiana* were quite similar, and ranged from 0.22-1.61 compared to the control gene *Nb_Actin3* (Fig. 1C).

The mRNA levels of the seven genes in *N. benthamiana* were higher than in *Arabidopsis*. The expression levels ranged from 136-127,535 folds (Figs. 1D, E, F). PLDα2 was expressed at an extremely low level in *Arabidopsis* leaves, making it difficult to directly amplify its CDS from cDNAs for cloning and transient expression.

We speculated that the position and number of introns within the *PLD*α*2* gene affected its transcription, mRNA output, and polyadenylation. In this way, depending on the position and number of introns, introns can even lead to degradation of gene expression. In addition, the expression of *PLD*α*2* genes may also require induction by corresponding exogenous pathogens. However, the transient expression of the gDNA of PLDα2 can increase its expression level 127,535 folds and facilitate the cloning of the CDS of PLDα2.

Our previous work has showed that PLDδ can negatively regulate the resistance to *pseudomonas syringae pv. maculicola 1* (RPM1) [14]. Under normal circumstances, the expression of the nucleotide-binding site leucine-rich repeat (NB-LRR) protein is very low in native plants. Next, we cloned the CDS of the NB-LRR gene from other species.

### 3.2 gDNAs from Oryza sativa can be transiently expressed in N. benthamiana leaves

Rice (*Oryza sativa*) is a staple crop and a model plant for research, purposes that we have used to characterize the NB-LRR gene *LOC_Os08g28460, LOC_Os08g28540*, and *LOC_Os08g10260* are three NB-LRR genes that were selected to study transient expression and splicing in *N. benthamiana* (Table 1). The YFP-HA tagged proteins LOC_Os08g28460 and LOC_Os08g28540 can be detected following transient expression in the leaves of *N. benthamiana*. The YFP-HA tagged LOC_Os08g10260 protein has a weaker but detectable level of protein expression (Fig. 3A). All three CDSs were amplified and sequenced to confirm the correct transcription and splicing in the transient expression system.

**Fig. 3.**
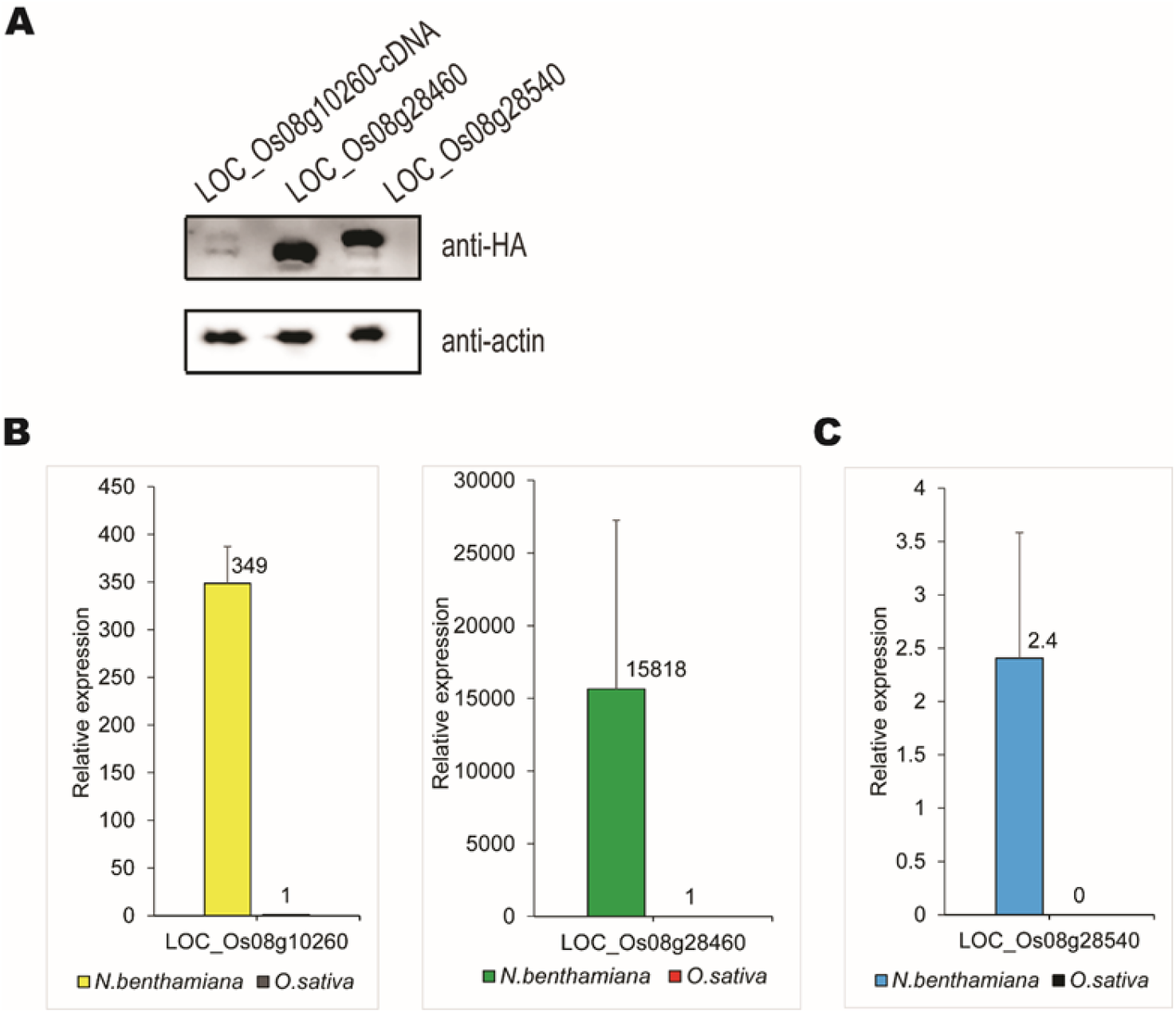
Transient expression of the gDNAs from *O. sativa* in leaves of *N. benthamiana* (A) The CDS of *LOC_Os08g10260, LOC_Os08g28460* and *LOC_Os08g28540* expressed YFP-HA tagged proteins in *N. benthamiana*. Expression constructs containing CDS or gDNAs were transiently expressed in *N. benthamiana* leaves. The CDS of *LOC_Os08g10260* was cloned from *N. benthamiana* leaves that expressed its gDNA. The total proteins were harvested and analyzed at 48 hpi. (B) Comparison of the relative expressions of *LOC_Os08g10260* and *LOC_Os08g28460* in *N. benthamiana* and *O. sativa*. The expression levels of *Os_Actin* (AB047313) and *Nb_Actin3* were used as internal controls. The relative expression level of each gene in *O. sativa* was set as 1. The values represent the average of three replicate experiments ± SD. (C) The relative expression of *LOC_Os08g28540* in *N. benthamiana* and in *O. sativa*.

The expression levels of the three genes were analyzed in rice and *N. benthamiana* by qRT-PCR. The mRNA levels of *LOC_Os08g10260* and *LOC_Os08g28460* in *N. benthamiana* were 349 folds and 15818 folds than in rice, respectively (Fig. 3B). The expression of *LOC_Os08g28540* was so low that the mRNA level could not be detected in the leaves of rice, whereas its mRNA level was 2.4 folds higher than the control gene *Nb_Actin3* in *N. benthamiana* (Fig. 3C).

### 3.3 Transient expression of gDNAs from other species in N. benthamiana

To further verify the application of the methods described above, we randomly selected NB-LRR genes from other species, including two dicots (*Brassica napus, Glycine max*) and two monocots (*Sorghum bicolor, Triticum aestivum*) (Table S2). Seven out of the nine NB-LRR genes from *Brassica napus* and three out of the five NB-LRR genes from *Glycine max* formed the predicted CDS. One out of the four NB-LRR genes from *Sorghum bicolor* and one out of two NB-LRR genes from *Triticum aestivum* formed the predicted CDS. As long as the predicted CDSs can be isolated, the expression of corresponding proteins can be detected in *N. benthamiana* (data not shown). The NB-LRR genes that did not form the predicted CDSs in *N. benthamiana* were cloned and sequenced from the leaves of the corresponding species, and compared to the sequences extracted from *N. benthamiana*. Interestingly, one or two introns were not spliced in the mRNA of *BnaA01g00570D, BnaA01g00670D*, and *Glyma*.*01G025400* in *N. benthamiana* according to the predicted CDS yet the sequencing results showed that the CDS of the three genes cloned from *N. benthamiana* were the same as those cloned from their species (Table S2). These results indicated that these forms are new spliced variants in these genes and that the splicing predictions for these genes need to be verified.

Overall, our results indicated that gDNAs from *Arabidopsis thaliana, Oryza sativa, Brassica napus*, and *Glycine max* can express correctly spliced mRNAs in a transient expression system of *N. benthamiana*, but gDNAs from *Sorghum bicolor* and *Triticum aestivum* do not function in this system.

### 3.4 The precursor messenger RNA of Arabidopsis PLDγ3 has spliced variants in N. benthamiana

There are two splice variants PLDγ3.1 and PLDγ3.2 in *Arabidopsis* (Fig. 4A). An extra 90 bp intron (548-637) was spliced out in PLDγ3.2 compared to the sequence of PLDγ3.1. These two variants can be determined with specific primers (Fig. 4B). We determined whether the gDNA of PLDγ3 can also form splice variants in *N. benthamiana*. Our results showed that these two splice variants may also be present in *N. benthamiana*. We also detected the ratio of the two variants by qRT-PCR. The ratio of PLDγ3.2 to PLDγ3.1 in the leaves of *N. benthamiana* was 1.24, whilst the ratio was 0.5 in the leaves of *Arabidopsis*, indicating that although *Arabidopsis* and *N. benthamiana* are both model plants, they have distinct functional differences (Fig. 4C).

**Fig. 4.**
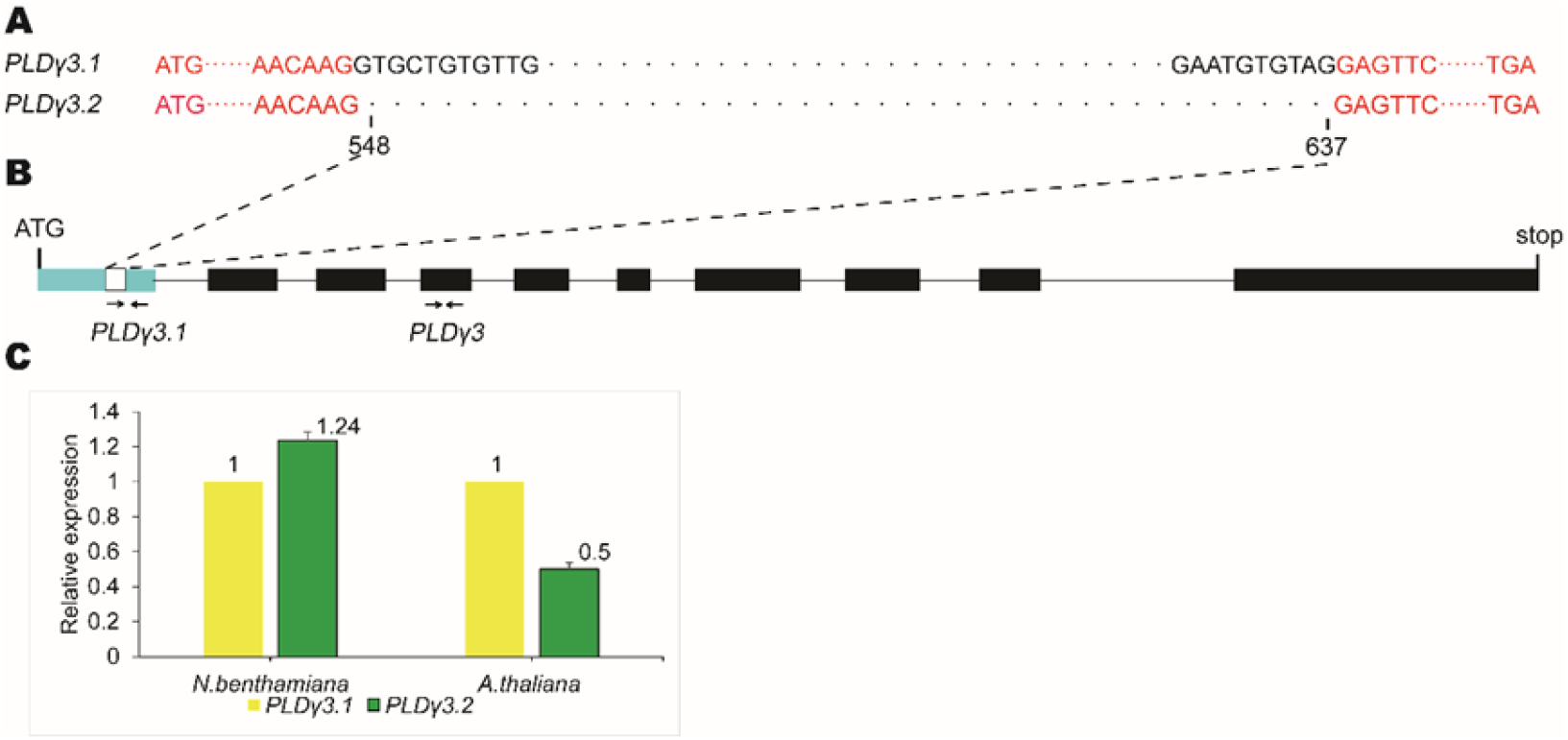
The mRNA of AtPLDγ3 has spliced variants in *N. benthamiana*. (A) Sequence alignment of PLDγ3.1 and PLDγ3.2. Two splice variants were determined from the sequencing of the CDS of AtPLDγ3 synthesized in *N. benthamiana*. PLDγ3.1 had an extra 90 bp intron in the range of 548 bp to 637 bp corresponding to the cDNA sequence of PLDγ3.2. (B) Two primer pairs are used to discriminate between PLDγ3 and PLDγ3.1. The arrows indicate the primers used to check the spliced variants. The PLDγ3 primer pair was used to analyze the total expression of PLDγ3.1 and PLDγ3.2, The PLDγ3.1 primer pair was used to analyze the expression of PLDγ3.1. (C) The relative expression of PLDγ3.1 and PLDγ3.2 in *Arabidopsis* and *N. benthamiana*. The relative expression level of PLDγ3.1 was set as 1. The values represent the average of three replicate experiments ± SD.

## 4. Discussion

In this study, we demonstrate an attractive alternative approach for genes with CDS that are difficult to clone using conventional methods. Our approach has several distinct advantages over established systems. gDNA can be easily amplified from plant genomic DNA, CDS are usually cloned from cDNA but are dependent on the presence and abundance of the target mRNA in the extracted plant tissues. This process is challenging for genes expressed at extremely low levels or have spatial and/or temporal expression patterns in native plants (e.g. NB-LRR genes). However, the efficiency of expressing proteins with gDNA may be lower compared to CDS as the synthesis of the LOC_Os08g10260 protein cannot be expressed with gDNA, but can be detected when expressed with the CDS.

Also, the high abundance of target mRNAs can be synthesized in the leaves of *N. benthamiana*. The levels and patterns of gene expression are controlled by the native promoters in plant genes. Genes such as *PLD*β*2* in *Arabidopsis*, and the NB-LRR gene *LOC_Os08g28540* in *Oryza sativa* are expressed at extremely low expression levels, making it difficult to clone CDSs amplified from cDNAs for cloning and transient expression. In this study, we used the constitutive CaMV 35S promoter to express the target genes in *N. benthamiana*, and the relative expression of the transiently expressed genes increased significantly. For example, the relative expression of *PLD*β*2* increased 65,000 folds, and there was no detectable expression of the NB-LRR gene *LOC_Os08g28540* in leaves of *Oryza sativa*. In comparison, the expression level of *LOC_Os08g28540* was 2.4 folds higher than the internal control *Nb_Actin3* in *N. benthamiana*.

Genes from three dicots (*Arabidopsis, Brassica napus*, and *Glycine max*) and three monocots (*Oryza sativa, Sorghum bicolor* and *Triticum aestivum*) were chosen to determine the accuracy of the CDSs cloned with *N. benthamiana*. Our results showed that this method is suitable for genes from *Arabidopsis thaliana, Oryza sativa, Brassica napus*, and *Glycine max*, but not for genes from *Triticum aestivum* and *Sorghum bicolor*. These data may be explained by several reasons. Firstly, the small number of candidate NB-LRR genes can result in statistical challenges. Secondly, the splice variants of certain mRNAs are tissue specific. Since the genes are expressed in *N. benthamiana* leaves, the leaf specific splice variant should be dominant. Thirdly, the expression of NB-LRR genes may be dependent on specific environmental or inducing factors or only when corresponding effectors are detected. Finally, the two species that were found to have limited efficacy were monocots plants, *N. benthamiana* is a dicot plant that is evolutionary different to monocot plants. For example, coiled-coil (CC)-NB-LRR and toll/interleukin 1 receptor (TIR)-NB-LRR genes both exist in *Arabidopsis*, yet only CC-NB-LRR genes are present in *Oryza sativa*, and so only some NB-LRR genes can be correctly expressed across species [15].

## 5. Conclusion

In short, we aimed to determine if plant gDNAs can be used for transient expression. We demonstrated that gDNAs from *Arabidopsis thaliana, Oryza sativa, Brassica napus*, and *Glycine max* can be correctly transcribed and spliced into mRNAs in *N. benthamiana*, whilst gDNAs from *Triticum aestivum* and *Sorghum bicolor* are not effective. In conclusion, our results suggested that the transient expression system of *N. benthamiana* is a useful tool for the cloning of CDSs of plant genes.

## Supplementary materials

**Table S1** *Selected genes from Arabidopsis and Oryza sativa*.

**Table S2** *Selected genes from other dicot and monocot plants*.

## References

[1] J. Reed, A. Osbourn, Engineering terpenoid production through transient expression in Nicotiana benthamiana, Plant Cell Reports, 37 (2018) 1431–1441. DOI: 10.1007/s00299-018-2296-3.

[2] S. Nandi, A.T. Kwong, B.R. Holtz, R.L. Erwin, S. Marcel, K.A. McDonald, Techno-economic analysis of a transient plant-based platform for monoclonal antibody production, mAbs, 8 (2016) 1456–1466. DOI: 10.1080/19420862.2016.1227901.

[3] F. Le Mauff, C. Loutelier□Bourhis, M. Bardor, C. Berard, A. Doucet, M.A. D’Aoust, L.P. Vezina, A. Driouich, M.M.J. Couture, P. Lerouge, Cell wall biochemical alterations during Agrobacterium□mediated expression of haemagglutinin□based influenza virus□like vaccine particles in tobacco, Plant Biotechnology Journal, 15 (2017) 285–296. DOI: 10.1111/pbi.12607.

[4] S. Chebolu, H. Daniell, Chloroplast-derived vaccine antigens and biopharmaceuticals: expression, folding, assembly and functionality, Current topics in microbiology and immunology, 332 (2009) 33–54. DOI: 10.1007/978-3-540-70868-1_3.

[5] T.V. Komarova, S. Baschieri, M. Donini, C. Marusic, E. Benvenuto, Y.L. Dorokhov, Transient expression systems for plant-derived biopharmaceuticals, Expert review of vaccines, 9 (2010) 859–876. DOI: 10.1586/erv.10.85.

[6] J. Huang, P. Jia, X. Zhong, X. Guan, H. Zhang, Z. Gao, Ectopic expression of the Arabidopsis mutant L3 NB-LRR receptor gene in Nicotiana benthamiana cells leads to cell death, Gene, 906 (2024). DOI: 10.1016/j.gene.2024.148256.

[7] M. Walter, C. Chaban, K. Schütze, O. Batistic, K. Weckermann, C. Näke, D. Blazevic, C. Grefen, K. Schumacher, C. Oecking, K. Harter, J. Kudla, Visualization of protein interactions in living plant cells using bimolecular fluorescence complementation, The Plant Journal, 40 (2004) 428–438. DOI: 10.1111/j.1365-313X.2004.02219.x.

[8] M.D. Edge, A.R. Greene, G.R. Heathcliffe, P.A. Meacock, W. Schuch, D.B. Scanlon, T.C. Atkinson, C.R. Newton, A.F. Markham, Total synthesis of a human leukocyte interferon gene, Nature, 292 (1981) 756–762. DOI: 10.1038/292756a0.

[9] X. Wu, X. An, J. Lu, J.-d. Huang, B. Zhang, D. Liu, X. Zhang, J. Chen, Y. Zhou, Y. Tong, Rapid Assembly of Multiple-Exon cDNA Directly from Genomic DNA, PLoS ONE, 2 (2007). DOI: 10.1371/journal.pone.0001179.

[10] Z. Wang, X. Liu, J. Yu, S. Yin, W. Cai, N.H. Kim, F. El Kasmi, J.L. Dangl, L. Wan, Plasma membrane association and resistosome formation of plant helper immune receptors, Proceedings of the National Academy of Sciences, 120 (2023). DOI: 10.1073/pnas.2222036120.

[11] M. Adnan, G. Morton, S. Hadi, Analysis of rpoS and bolA gene expression under various stress-induced environments in planktonic and biofilm phase using 2−ΔΔCT method, Molecular and Cellular Biochemistry, 357 (2011) 275–282. DOI: 10.1007/s11010-011-0898-y.

[12] L. Fan, S. Zheng, D. Cui, X. Wang, Subcellular distribution and tissue expression of phospholipase Dalpha, Dbeta, and Dgamma in Arabidopsis, Plant Physiol, 119 (1999) 1371–1378. DOI: 10.1104/pp.119.4.1371.

[13] C. Wang, X. Wang, A Novel Phospholipase D of Arabidopsis That Is Activated by Oleic Acid and Associated with the Plasma Membrane, Plant Physiology, 127 (2001) 1102–1112.

[14] X. Yuan, Z. Wang, J. Huang, H. Xuan, Z. Gao, Phospholipidase Dδ Negatively Regulates the Function of Resistance to Pseudomonas syringae pv. Maculicola 1 (RPM1), Frontiers in plant science, 9 (2019). DOI: 10.3389/fpls.2018.01991.

[15] J.D.G. Jones, R.E. Vance, J.L. Dangl, Intracellular innate immune surveillance devices in plants and animals, Science, 354 (2016) aaf6395. DOI: 10.1126/science.aaf6395.

