## Supplementary Table 1 for "The application of *Nicotiana benthamiana* as a Transient Expression Host to Clone the Coding Sequences of Plant Genes"

**Table S1** *Selected genes from Arabidopsis and Oryza sativa.*

| Gene name  (Gene locus) | Length of gDNA (bp) | Length of CDS (bp) | Intron numbers |
| --- | --- | --- | --- |
| *PLDα2 (AT1G52570)* | 2612 | 2433 | 2 |
| *PLDβ1 (AT2G42010)* | 4973 | 3252 | 9 |
| *PLDβ2 (AT4G00240)* | 4339 | 2784 | 9 |
| *PLDδ (AT4G35790)* | 4102 | 2574 | 9 |
| *PLDε (AT1G55180)* | 2573 | 2289 | 3 |
| *PLDγ1 (AT4G11850)* | 3586 | 2577 | 9 |
| *PLDγ3 (AT4G11840)* | 3731 | 2511 | 9 |
| *LOC_Os08g10260* | 3438 | 2985 | 1 |
| *LOC_Os08g28460* | 4676 | 2214 | 2 |
| *LOC_Os08g28540* | 4044 | 2730 | 1 |
