## Supplementary Table 2 for "The application of *Nicotiana benthamiana* as a Transient Expression Host to Clone the Coding Sequences of Plant Genes"

**Table S2** *Selected genes from other dicot and monocot plants.*

| Tested species | Genes shown predicted splicing (intron numbers) | Genes shown different splicing (not removed introns/total introns) |
| --- | --- | --- |
| Brassica napus | *BnaA01g00520D* (1)  *BnaA01g03030D* (1)  *BnaA01g02960D* (2)  *BnaA01g00220D* (4)  *BnaA03g34210D* (2)  *BnaA03g34370D* (3)  *BnaA03g30890D* (2) | *BnaA01g00570D^a^* (1/2)  *BnaA01g00670D^a^* (2/2) |
| *Glycine max* | *Glyma.01G032600* (6)  *Glyma.01G027000* (1)  *Glyma.01G038600* (2) | Glyma.01G025400^a^ (1/1)  *Glyma.01G030100^c^* (2/2) |
| *Triticum aestivum* | *Traes_1AL_5B26F6A14* (1) | *Traes_1AL_2AC682D6D^b^* (1/3) |
| *Sorghum bicolor* | *Sobic.001G000900* (1) | *Sobic.001G001000^c^* (3/3)  *Sobic.001G002100^c^* (3/3)  *Sobic.001G005900^b^* (3/4) |

***^a^*** *Sequencing results proved that the CDS cloned from N. benthamiana is the same as cloned from the corresponding native plant, although they are different according to the predicted CDS sequence (https://phytozome.jgi.doe.gov/pz/portal.html).*

***^b^*** *Sequencing results proved that the CDS cloned from N. benthamiana is different from the predicted CDS.*

***^c^*** *The CDS can’t be cloned from leaves of corresponding plant. The difference can’t be verified.*
